# Anodal M1 tDCS enhances online learning of rhythmic timing videogame skill

**DOI:** 10.1101/2023.11.22.568299

**Authors:** Anthony W. Meek, Davin Greenwell, Brach Poston, Zachary A. Riley

## Abstract

Transcranial direct current stimulation (tDCS) has been shown to modify excitability of the primary motor cortex (M1) and influence online motor learning. However, research on the effects of tDCS on motor learning has focused predominantly on simplified motor tasks. The purpose of the present study was to investigate whether anodal stimulation of M1 over a single session of practice influences online learning of a relatively complex rhythmic timing video game. Fifty-eight healthy young adults were randomized to either a-tDCS or SHAM conditions and performed 2 familiarization blocks, a 20-minute 5 block practice period while receiving their assigned stimulation, and a post-test block with their non-dominant hand. To assess performance, a performance index was calculated that incorporated timing accuracy elements and incorrect key inputs. The results showed that M1 a-tDCS enhanced the learning of the video game based skill more than SHAM stimulation during practice, as well as overall learning at the post-test. These results provide evidence that M1 a-tDCS can enhance acquisition of skills where quality or success of performance depends on optimized timing between component motions of the skill, which could have implications for the application of tDCS in many real-world contexts.

## Introduction

Motor learning is fundamental in everyday life for acquiring and honing skills that range in complexity from relatively simple, like reaching and pressing a button, to more complex skills requiring coordinated sequential actions such as learning a piece of music on an instrument or throwing a ball. Motor skill improvements can accrue during a single practice session (online) or after practice is completed (offline), with both contributing over time to long-term retention [1]. The online and offline skill changes comprise fast and slow stages of learning, with fast stage learning occurring early on, markedly during skill acquisition, and slow stage occurring later with incremental gains over multiple practice sessions [2, 3]. The motor learning process is underpinned by neuroplastic changes across a spatially distributed network of interconnected brain regions [4]. Which areas are involved and the extent to which plasticity within them may subserve learning, largely depends upon the characteristics of task and the stage learning [5].

Evidence of learning related cortical plasticity in the primary motor cortex (M1) after skill practice has indicated that it contributes to motor skill acquisition. M1 is crucial for the use-dependent acquisition and storage of muscle activation for fast and precise motions associated with skillful performance [6, 7]. When a motor skill is practiced, somatotopically specific changes arise via LTP-like plasticity within M1 that improve synaptic connectivity and efficiency amongst the ensemble of cells activated to generate the movements of the skill [8, 9]. The principal result of this process is enlarged cortical representations of the muscles involved in the task and/or facilitation of motor evoked potentials (MEPs), indicative of increased cortical excitability, as assessed by TMS [10, 11]. Short bouts of skill practice (10 to 30 minutes) have been shown to elicit acute increases in M1 excitability [11, 12]. Although early practice-related plasticity is relatively transient in the short term [13], it is believed to be an important initial step of fast motor learning and skill acquisition [2, 11]. Based on these short term mechanisms as well as the more permanent structural and functional cortical reorganization associated with extensive practice and skill expertise [10, 14–16], M1 is a site of particular interest for investigating motor learning.

Transcranial direct current stimulation (tDCS) is a non-invasive brain stimulation technique for modulating cortical excitability. By passing a current between two electrodes placed on the scalp, tDCS provides a sub-threshold, transcranial stimulation to the underlying cortical surface. Directional flow of the current between these electrodes determines the stimulation polarity (positive, Anodal; negative, Cathodal) [17]. Anodal tDCS (a-tDCS) in particular has been used to enhance cortical excitability and LTP-like plasticity in M1 corresponding with improved motor function [18–20]. tDCS has been shown to be particularly relevant in the acquisition and early consolidation phase of motor learning and evidence suggests that it is safe for facilitating motor learning in healthy individuals [20, 21] as well as those suffering from neurological disorders [22, 23].

Improved performance from tDCS application to M1 during practice has been reported for several upper limb motor tasks where skill is usually assessed through changes in speed (reaction or movement time), accuracy, the relationship between those two qualities, or reduced variability [19]. Most studies have used laboratory based tasks, such as the serial reaction time task (SRTT), sequential finger tapping task (SFTT), and sequential visual isometric pinch force task (SVIPT) that offer highly controlled, simplified environments [19, 20, 24]. Although lab-based tasks provide precise measurements and high degrees of control, they lack the complexity and visual-motor demands that most normal everyday tasks often involve, making generalization to real-world tasks limited. Nevertheless, these studies lend support to M1’s important role in fast stage learning and potential as a target for tDCS to benefit learning.

The benefit of M1 tDCS on motor skill acquisition has been demonstrated with an increasing number of tasks with expanding complexity [25–28]. Videogames are emerging as a potentially useful tool to study motor learning since they involve diverse combinations of perceptual, attentional, cognitive, and motor skills, and practice can lead to training-induced learning [29, 30]. Although there are reports of competitive gamers using tDCS to enhance performance [31], the effect of M1 tDCS in the context of gaming-based motor skill has only recently been initially investigated [26]. With comparable motor demands to laboratory tasks, but more visually complex environments and precise temporally-constrained inputs, video games present a novel way to study the effects of M1 tDCS on motor learning. Given the task dependency of tDCS effects for relatively simple tasks [19, 32, 33], it is unclear how M1 tDCS will influence the acquisition of a rhythmic sequence tapping task with complex visuomotor and auditory processing demands. Therefore, the purpose of this study was to investigate how M1 a-tDCS influences the acquisition of a timing-based video game skill.

## Methods

### Participants

58 total subjects (*n* = 29 per group; age 22.27 ± 2.78 yrs) participated in the study between the dates 2/13/2018 and 4/12/2018. Participants were free of neurological or musculoskeletal impairments that could impact performance of the task. Participants that were taking medications that could influence learning, such as stimulants for ADHD, were also excluded. Handedness was determined with the Edinburgh Handedness Inventory (EHI) [34]. Right-hand dominance was identified with scores > +40, left-hand dominance with scores < -40, while scores ≥-40 and ≤+40 were ambidextrous. In cases of ambidextrousness, the subject’s preferred writing hand was considered their dominant.

The study was a randomized, between-subjects, SHAM controlled design. Participants were randomized to an anodal tDCS group (a-tDCS) or SHAM stimulation group (SHAM) as they were screened into the study and were blinded to their condition for the duration of testing. The two groups were relatively matched for age (a-tDCS = 22.70 ± 2.96 yrs and SHAM = 22.34 ± 2.64 yrs) and sex (a-tDCS = 14 males, 15 females; SHAM = 15 males, 14 females). For handedness, considering preferred writing hand in the case of an ambidextrous inventory result, each group had 25 right-handed individuals and 4 left-handed individuals. All procedures were approved by Indiana University’s IRB and conducted according to the Declaration of Helsinki and all subjects provided their written informed consent before participation in the study.

### Procedures

Participants completed a single testing session while receiving a-tDCS or SHAM stimulation over M1 contralateral to the non-dominant (active) hand. During the session, subjects performed a timing-based dexterous video game task with the non-dominant hand. The protocol included 2 familiarization pre-test blocks, 5 practice blocks while receiving either a-tDCS or SHAM, and a post-test block 5 min after cessation of practice. Familiarization pre-test scores were used to determine whether the task difficulty was adequate to demonstrate motor learning during the practice period. If the task was too easy or too hard based on the familiarization trial performance, the difficulty was increased or decreased accordingly in the video game settings, and one additional familiarization trial was given at the new speed (Fig. 1B).

**Fig 1.**
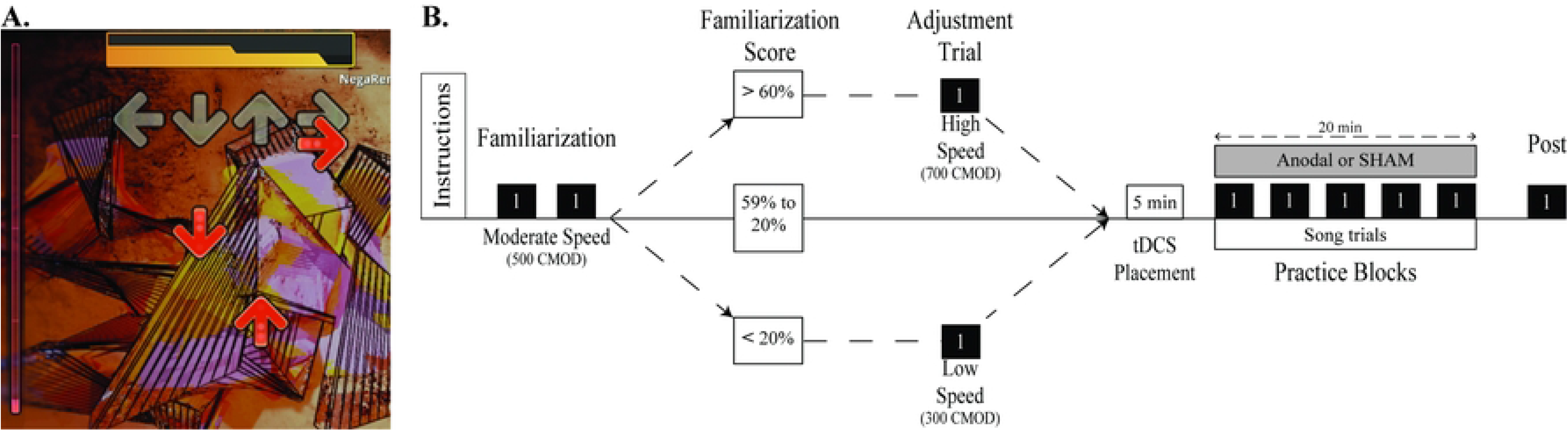
Gameplay and experimental procedures timeline. (a) Screen shot of Step Mania gameplay showing scrolling arrow icons and stationary arrow silhouettes. (b) Study procedures timeline.

### Step Mania task

Step Mania (https://www.stepmania.com/) is an open-source videogame where sequences of directional arrow icons (up, down, left, right) scroll upwards toward stationary arrow silhouettes (Fig. 1A) and the objective is to press the appropriate arrow keys whenever a corresponding scrolling icon is perfectly aligned with (i.e. centered on) its silhouette.

Participants sat at a desk with a keyboard positioned with the arrow keys in front of the non-dominant hand. Subjects were instructed to only use one hand to tap the keys and to only tap keys one time per cue. However, no explicit instructions were given assigning specific digits to particular keys. A brief demonstration of the gameplay was shown to the participants as the objective was explained (*i.e.* press correct keys with optimal timing and avoid unnecessary/incorrect key strokes).

The same 219 cue pattern was used for all test blocks. One of the 219 cues required the left and right arrow key to be struck simultaneously, thus the pattern consisted of 55 up, 56 down, 55 left and 54 right key strokes (220 total key taps). This could not be changed through the available settings. Each testing block comprised one complete pattern. Each keystroke was categorized relative to a time window centered (time = 0s) on perfect overlap of the scrolling icon and its stationary silhouette. These timing windows were built into the program as flawless (0 to ± 0.0225s), perfect (±0.0225 to ±0.045s), great (±0.045 to ±0.090s), good (±0.090 to ±0.135s), boo (±0,135s to ± 0.180s), and miss (> ± 0.180s). To note, the first two time windows (flawless and perfect) were 22.5ms, while the other windows were 45ms.

### Baseline assessment and testing

Subjects performed 2 familiarization trials with 500 continuous modifier (CMOD) difficulty (sets the arrow scroll speed; “arrow heights” moved per minute). An initial proficiency score was calculated (*Eq. 1*) indicative of the number of cues being met with a tap within scoring distance. This allowed for difficulty adjustment to ensure that the task was appropriately difficult for performance gains to be demonstrated within a single practice session. It was determined in pilot testing that if too many cues were entirely missed or nearly unscored based on the inbuilt timing windows, the task overwhelmed participants and they struggled to improve. Conversely, individuals who, in their familiarization trials demonstrated very precise timing and few misses, had little room for growth.

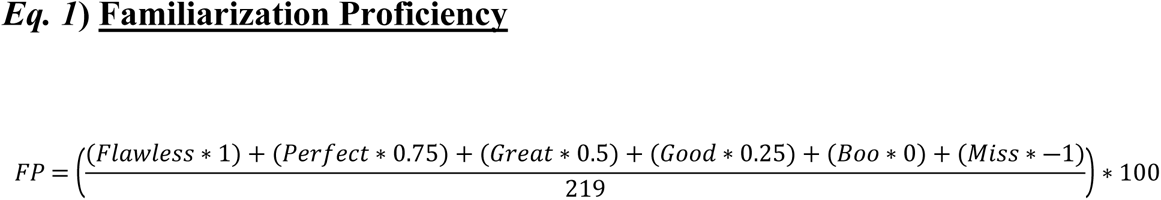

For familiarization proficiency scores between 20 and 60, the default 500 CMOD was maintained throughout testing. For scores > 60 or < 20, the scrolling speed was adjusted up or down by 200, respectively (Fig. 1B). The music tempo (210 bpm) was not affected by the CMOD change, it only affected the cue scrolling speed. Skill-adjusted subjects performed an additional familiarization block at the new speed that was used for all subsequent blocks. The last familiarization block was used as the pre-test measure for all subjects.

After familiarization and pre-test, tDCS electrodes were positioned during a 5 min break. Subjects then completed a 20 min practice period while they received either a-tDCS or SHAM during 5 practice blocks with 2 minutes of rest between blocks. The stimulation electrodes were then removed and subjects completed a post-test trial 5 minutes after completing practice.

### Evaluating performance

At the end of each trial, the game provides feedback in the form of a numerical “game score” as well as an aggregate overview of the categorization of each keystroke timing (i.e. miss, boo, good, great, perfect, and flawless). However, this fails to account for any excess or aberrant keystrokes that fall outside of the inbuilt timing windows. For this reason, we opted to collect additional data using a separate keystroke logging software. This allowed us to devise an alternate scoring mechanism, or performance index (PI), that better reflects changes in task performance.

Two aspects of Step Mania performance are related to skill – hitting only the correct inputs and hitting them at the correct time. Both are interrelated as qualities of the task, but a change in one does not necessarily precipitate a proportional change in the other. With the guiding premise that newly acquired movement sequences are segmented, inaccurate, and jerky, whereas learned sequences are cohesive, accurate and smooth [6], a PI was calculated to incorporate incorrect inputs (Key Error Rate, KER) and temporal accuracy (2 values: Temporal Accuracy, TA; Tap Distribution Ratio, TDR) into a single value to reflect the skill associated with a certain quality of performance:

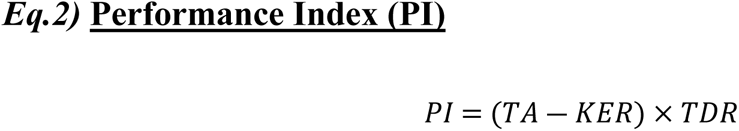

A description is provided below for each of the constituent variables in the PI equation and how they are derived from the data provided by the game. Example data are shown after each equation to illustrate each step of the process.

### Temporal Accuracy (TA)

TA was quantified by assigning point values to the timing windows, so that “flawless” keystrokes awarded 1.0 pt and each subsequent window awarded 0.2 pts less, resulting in 0 pts for each “miss”. TA provided a base value reflecting temporal accuracy of the entire 220 constituent inputs of each trial.

This is different from the Familiarization Proficiency equation we used to evaluate initial ability because we did not want to penalize a “miss” with a negative score and instead just not award any points in that case. This distinction was made because in the initial familiarization, if the speed of the game was too fast for the subject they would get overwhelmed and miss several arrows in a row without even attempting any inputs, in which case a measure was needed to clearly inform us to slow the game down. In the case of the timing accuracy measure (*Eq. 2*), the subject would typically only miss an arrow occasionally and therefore just not be awarded points for it.

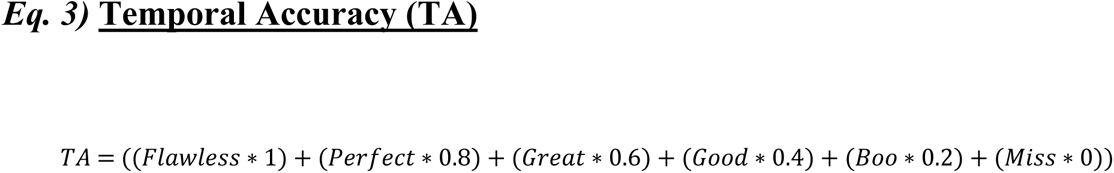

For a subject with 72 Flawless, 69 Perfect, 58 Great, 10 good, 3 Boo and 7 miss:

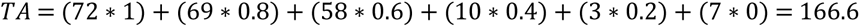

### Key Error Rate (KER)

KER evaluated execution errors such as pressing multiple keys simultaneously, tapping the same key several times, or pressing a cycle of keys ‘searching’ for the correct one. Imprecise and unstable finger movements relate to learning in aspects of the skill, irrespective of timing (i.e. key sequence order, hand positioning, etc). Utilizing the extra keystroke data collected during the game, the differences between actual key stroke totals and the number of times the arrow keys appear in the sequence were calculated for each trial and used in Eq. 3 to produce a scaled error value relative to the total number of cues.

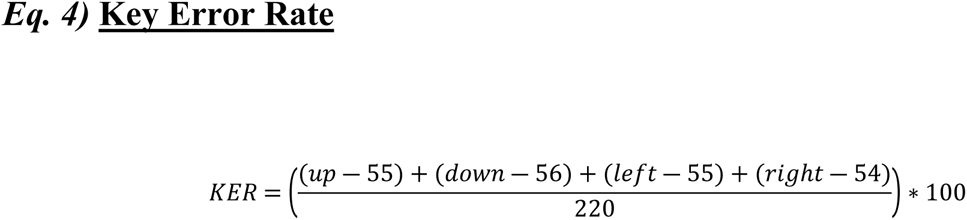

For a subject with 56 up, 57 down, 53 left and 53 right arrow:

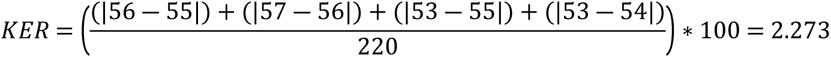

### Tap Distribution Ratio (TDR)

In rhythm gaming, ratios of the tap totals in the different timing windows are commonly used in conjunction with timing scores because, as a consequence of binning taps into scored timing windows that are relatively wide, attempts can yield arbitrarily similar TAs despite the distribution of taps in the windows reflecting different levels of play. We adapted this strategy to further differentiate skill in the task and enhance sensitivity in the measure to changes in the quality of performance. TDR adjusts scores based on the concentration of taps in the two best and two worst categories, with additional weighting based on the proportion of those that are in the best (Flawless; most skillful) and worst (Miss; least skillful) timing windows. To properly illustrate the function of this variable, the original TA data from above will be used as well as a second example that produces an identical 166.6 TA value.

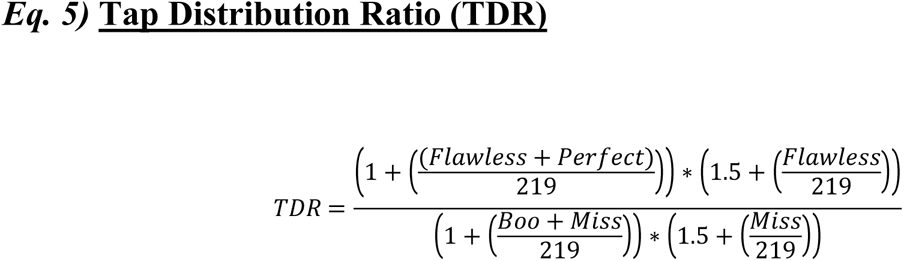

For a subject with 72 Flawless, 69 Perfect, 58 Great, 10 good, 3 Boo and 7 Miss:

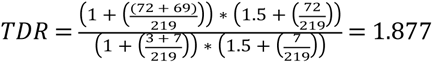

For a second subject with 82 Flawless, 46 Perfect, 67 Great, 18 Good, 2 Boo, 4 Miss:

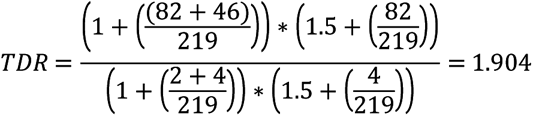

Assuming a similar KER value for both of these trials, the PI for each would be:

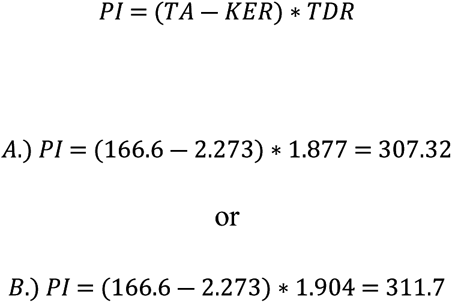

In utilizing a TDR factor to modify PI scores, we are able to better reflect the more subtle changes in subject performance that might otherwise be missed when relying on TA and KER values alone.

### tDCS

The location of M1 was approximated using the BA9 BA8 BA42 Location System [35] and a marker was used to denote this location on the scalp. A Soterix Medical 1×1 Low Intensity transcranial DC stimulator was used to deliver a-tDCS with parameters previously determined to be effective and safe (duration 20 min; current 1mA; active electrode over the marked M1 contralateral to the non-dominant hand and reference electrode over the ipsilateral supraorbital). Current was delivered through 5cm x 5cm (25cm^2^) rubber electrodes inside of saline-soaked sponges affixed to the head with rubber straps. For SHAM, the current was ramped up and down over 30s.

### Data Analysis and Statistics

A PI value was calculated for each trial. PI gain scores (block PI – baseline PI) were calculated for each practice block and the post-test to demonstrate the change in performance over time relative to baseline performance. Normality was tested with a Shapiro-Wilk test. Baseline PI scores were compared between groups with an independent *t*-test to test for baseline differences before gain scores were calculated.

Separate comparisons were performed for the practice blocks and the post-test block. A mixed ANOVA (2 groups x 5 blocks) with repeated measures on block was used to compare the practice gain scores and effect size were calculated as partial eta squared (η_p_ ^2^). For the post-test block, an independent *t*-test was used to compare the gain scores between the conditions and effect size was calculated as Cohen’s *d*. A p-value of 0.05 was considered statistically significant. Bonferroni post-hoc tests were used to determine differences for multiple comparisons. Data are presented as mean ± standard deviation in text and as mean ± standard error in figures. All data analysis was performed with SPSS 24.

## RESULTS

### Practice Blocks

Over the course of practice (B1 through B5), both groups gradually improved TA (range: 147.78 to 160.72) and TDR also modestly increased (range: 1.56 to 1.80). The rate of errors (KER) also improved (i.e. reduced) across practice for both groups (range: 3.83 to 2.71) (see Table 1).The calculated PI score for each block incorporated all 3 of these metrics.

**Table 1.** Mean TA, KER, and TDR values for each test block.

|  | Baseline | Practice |  |  |  |  | Post-Test |
| --- | --- | --- | --- | --- | --- | --- | --- |
|  |  | B1 | B2 | B3 | B4 | B5 |  |
| **a-tDCS** |  |  |  |  |  |  |  |
| TA | 135.66(19.81) | 149.84(19.16) | 152.85(20.02) | 154.31(15.99) | 159.42(15.04) | 160.72(16.01) | 166.63(15.57) |
| KER | 5.36(4.53) | 3.83(3.42) | 3.30(2.59) | 3.15(2.06) | 3.26(2.33) | 2.82(1.91) | 2.63(1.93) |
| TDR | 1.35(0.29) | 1.59(0.32) | 1.65(0.34) | 1.66 (0.28) | 1.76(0.28) | 1.80(0.31) | 1.92(0.31) |
| **SHAM** |  |  |  |  |  |  |  |
| TA | 134.96(27.41) | 147.78(22.42) | 150.78(22.32) | 154.21(19.79) | 152.66(17.71) | 155.36(19.73) | 159.62(18.98) |
| KER | 3.82(2.99) | 3.32(1.97) | 2.74(2.33) | 2.90(2.16) | 2.98(2.30) | 2.71(1.82) | 2.03(1.79) |
| TDR | 1.37(0.36) | 1.56(0.35) | 1.62(0.35) | 1.68(0.35) | 1.64 (0.32) | 1.69(0.37) | 1.78(0.37) |
| TA - Temporal Accuracy; KER - Key error rate; TDR - Tap Distribution Ratio; values are Mean (SD) |  |  |  |  |  |  |  |

TA - Temporal Accuracy; KER - Key error rate; TDR - Tap Distribution Ratio; values are Mean (SD) Practice block PI gain scores were calculated relative to baseline to evaluate how overall performance of the two groups changed during practice. An independent *t*-test on the baseline PI values showed a-tDCS (181.80 pts ± 66.04) and SHAM (188.97 pts ± 79.39) performance was similar prior to practice (*t*(56) = -0.37, *P* = 0.71). An a-tDCS subject was missing error data for two practice trials and thus was excluded from the practice block analysis (a-tDCS *n* = 28, SHAM *n* = 29).

A Shapiro-Wilk test showed normally distributed gain scores (*P* = 0.278 to 0.977) for all cells of the design except a-tDCS B1 (P = 0.004) and a-tDCS B4 (P = 0.01). Since ANOVA type 1 error rate is fairly robust to normality violations [36–38] the mixed ANOVA was still performed. Homogeneity of variances was confirmed by Levene’s test (*P* = 0.25 to 0.65). A Huynh-Feldt correction was used to adjust for a sphericity violation (*P* = 0.003). There was a statistically significant interaction between the stimulation group and block on practice gain scores (F[3.466, 190.605] = 3.042, *P* = 0.024, η_p_ ^2^ = 0.052). Post hoc analysis of the simple main effects for group revealed a significant difference between conditions at B4 (*P* = 0.008) and B5 (*P* = 0.046), where a-tDCS (B4: 100.02pts ± 47.21; B5: 109.05pts ± 60.92) resulted in greater PI improvement than SHAM (B4: 62.05pts ± 55.75; B5: 76.58pts ± 59.16). Post hoc analysis for block showed that B3 and B5 SHAM gain scores were significantly different from B1 (*P* = 0.002 and *P* = 0.005). For a-tDCS, B4 and B5 gain scores were statistically significantly higher than blocks 1 (both, *P* < 0.001), 2 (*P* = 0.049 and *P* = 0.008) and 3 (*P* = 0.005 and *P* = 0.002).

### Pre-test to Post-test

One SHAM subject was missing error data for the post-test and thus was excluded from analysis (a-tDCS *n* = 29, SHAM *n* = 28). Normality of the gain scores was confirmed with a Shapiro-Wilk test (*P* = 0.661 to 0.540). An independent *t*-test on the gain scores from baseline to post-test showed a statistically significant difference between conditions (*t*(55)= 2.234, *P* = 0.030, Cohen’s *d* = 0.592), with a-tDCS (137.844pts ± 66.218) leading to greater improvement compared to SHAM (99.501pts ± 63.28) (Fig. 3).

## DISCUSSION

These results indicate M1 a-tDCS has a beneficial effect on the acquisition of a dexterous timing-based video game compared to SHAM stimulation. Task improvement was indicated by higher gain scores, representing increased cumulative performance with better temporal accuracy and reduced error rate. Although both conditions demonstrated significant differences in performance across practice, the level of improvement with a-tDCS at B4 and B5 was significantly greater than that attained by SHAM (Fig. 2). Likewise, considering only the change in performance from baseline to post-test, a-tDCS during practice resulted in a greater overall change in PI compared to SHAM stimulation (Fig. 3).

**Fig 2.**
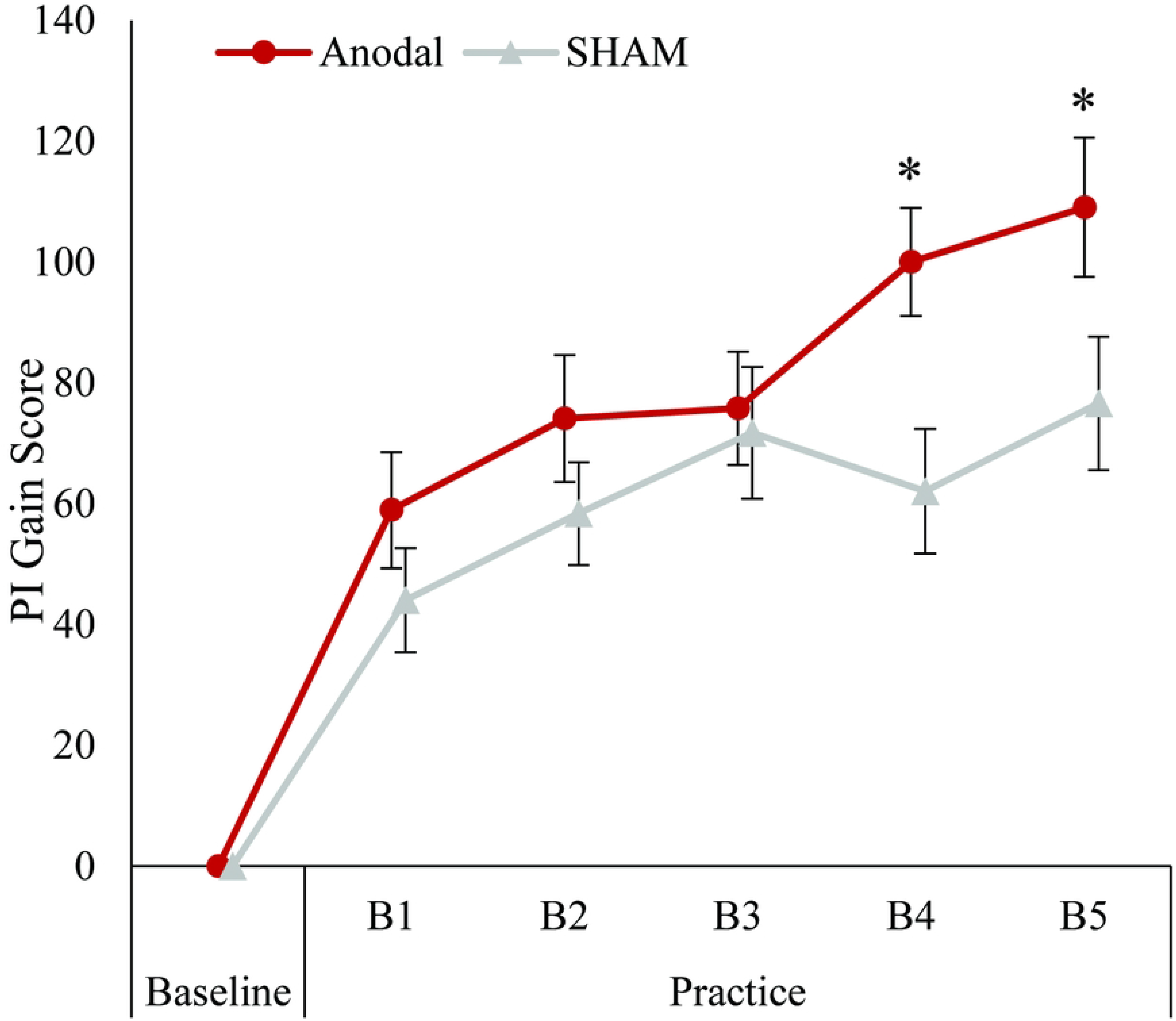
Practice block gain scores. Data show the practice block PI gain scores relative to baseline (i.e. last familiarization block) for a-tDCS (*n* = 28) and SHAM (*n* = 29) conditions. Whiskers denote the standard error. Both groups displayed significant increases in performance across practice. a-tDCS gain scores were significantly greater than SHAM at B4 and B5. *Indicates significant difference from SHAM (*P* < 0.05).

**Figure 3.**
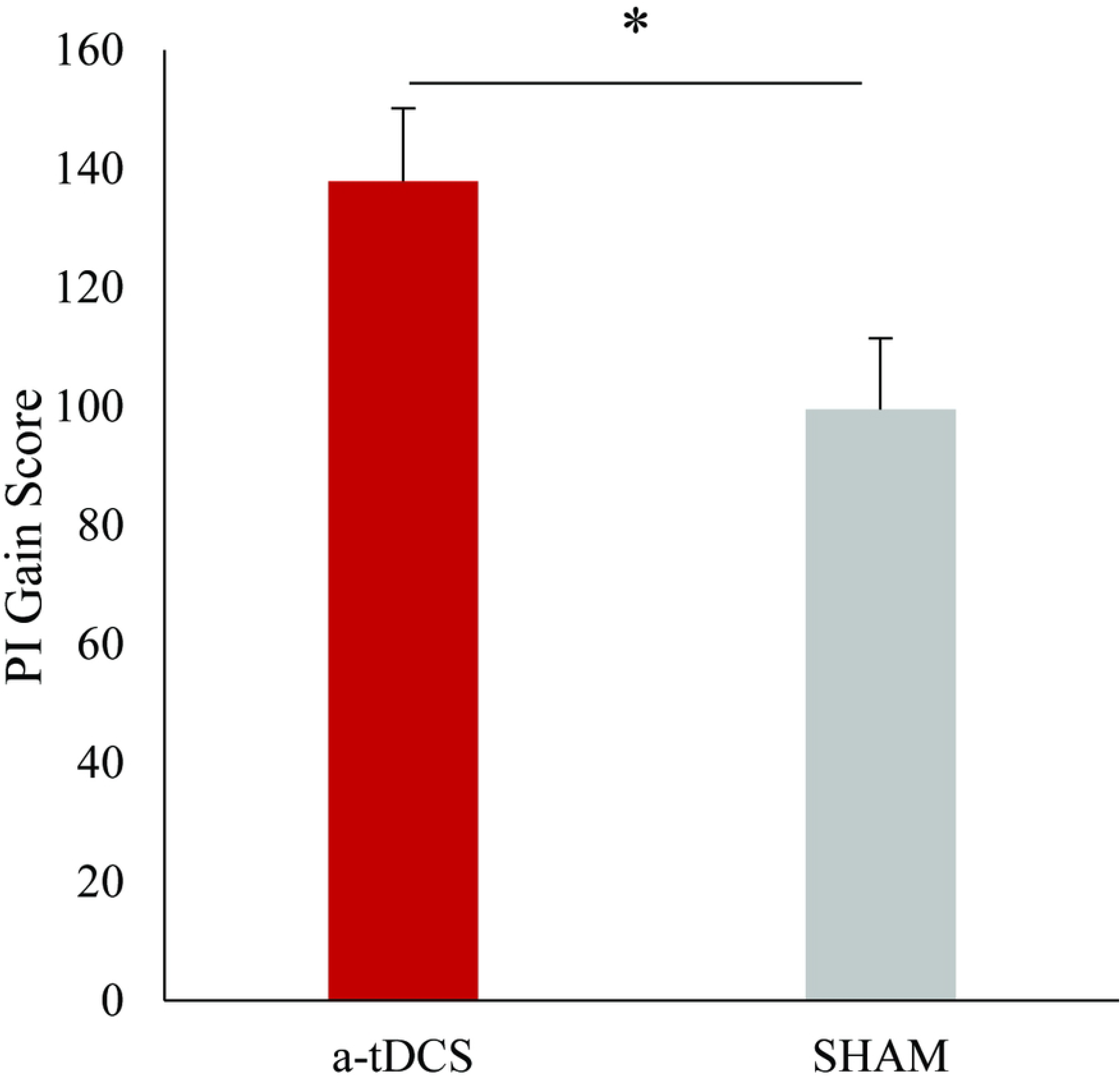
Overall gain scores. Data show the gain scores from the baseline to the post-test block for a-tDCS (*n* = 29) and SHAM (*n* = 28) conditions. Whiskers denote the standard error. *Significant difference between a-tDCS and SHAM (*P* < 0.05).

Our results align with previous findings that indicate M1 a-tDCS is a beneficial tool to enhance acquisition and performance of dexterous motor skills [19, 32, 39]. However, most of the previous evidence is based on simple laboratory tasks that do well isolating and measuring specific learning processes, but lack the variety of parameters many real-world motor skills involve. Step Mania is a complex rhythmic timing video game that provides ample explicit feedback. Even though M1 is important in skill acquisition, the effects of tDCS on motor learning are known to be task-dependent [32, 33] and when several areas outside M1 influence task performance, it has been suggested that M1 modulation alone may be insufficient to enhance motor learning [19]. Within this context, our finding that M1 was a viable target for tDCS to enhance acquisition and performance of a novel complex task are particularly interesting.

A potential issue with a global PI score is that it may indicate performance changes, but lack detail about how specific aspects of execution changed to improve performance. Partly due to our hypothesis that a-tDCS would enhance learning in a complicated task not singularly defined by one aspect of performance, but also due to game data extraction limitations, our methodology lacked the resolution and granularity typically needed to detect differences in individual aspects of performance. However, visual inspection of the general raw data trends (TA windows and KER) suggests the timing element of the task was the driving factor differentiating performance gains between the groups (Fig. 4 A-C). Both groups demonstrated rapid initial KER changes characteristic of sequence learning tasks [6] with similar trends across practice and post-test (Fig. 4C) indicating the groups acquired the sequence order similarly. Regarding timing quality, both a-tDCS and SHAM had more taps in the best TA window with a concurrent decrease in the worst window (Fig. 4B). By practice B5 and post-test, however, there is a clear TA advantage favoring a-tDCS with 10.3 and 13.3 more taps on average than SHAM in the best two timing windows combined and 2.9 and 2.7 fewer misses in B5 and the post-test, respectively (Fig. 4A).

**Fig 4.**
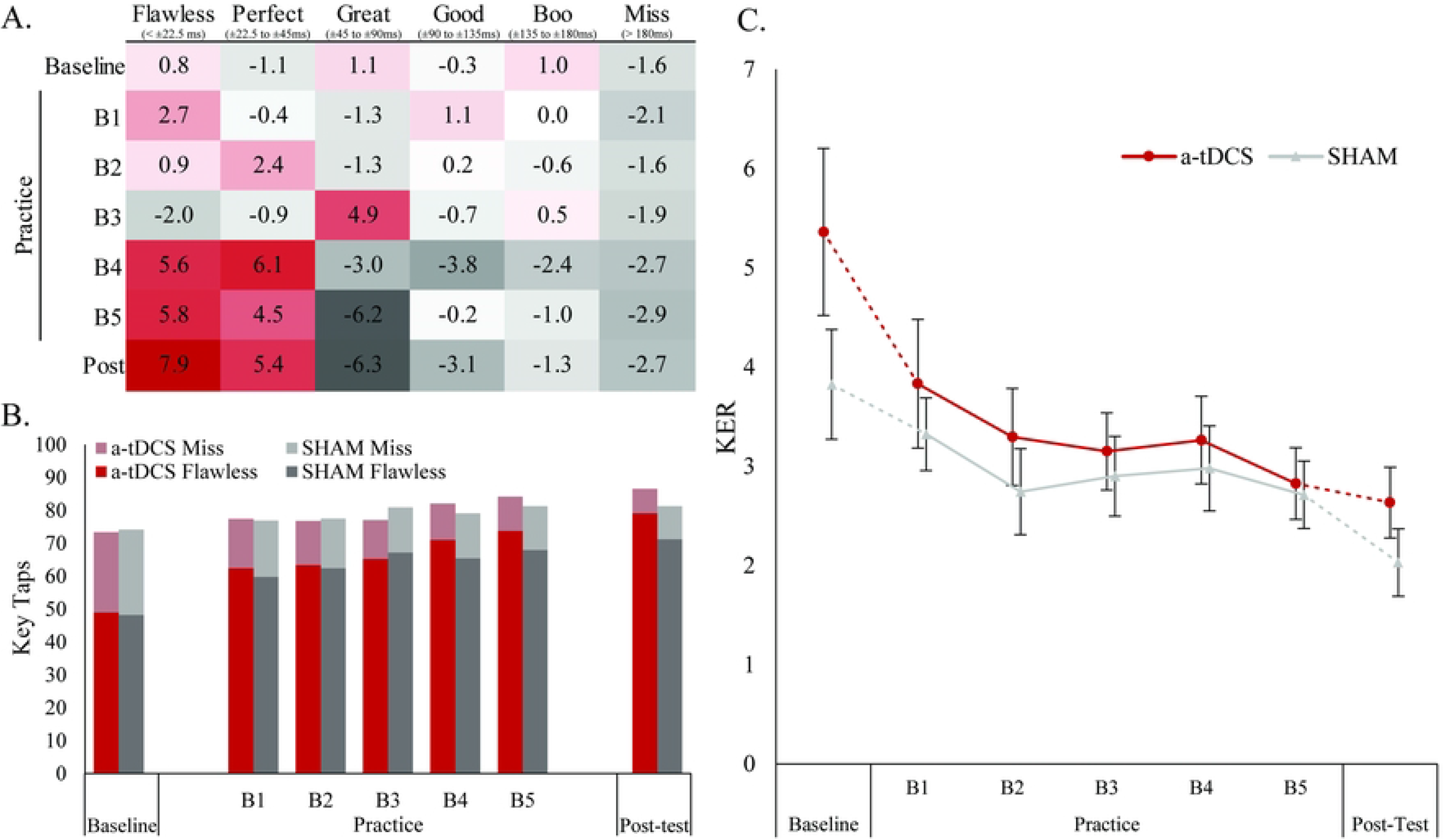
Timing window tap concentrations and key error rates. The color gradient table (A) shows the difference between the average a-tDCS tap distribution and average SHAM tap distribution (a-tDCS total – SHAM total) for each trial. Darker red tones signify differences favoring a-tDCS (positive values) and darker grey tones signify differences favoring SHAM (more negative). The stacked bar graph (B) shows the total number of taps in the best (“Flawless”, 0 to ± 0.0225s) and worst (“Miss”, > ± 0.180s) timing windows across trials. The total height of the bars represents the combined total number of taps falling within those two windows. The dark bottom portions of the bars indicate the proportion of flawless taps and the lighter top portions represent misses. The line graph (C) shows the KER trend from baseline, across practice to the post-test. Whiskers denote standard error.

It is likely that M1 tDCS facilitated chunking of Step Mania subsequences as previous work has demonstrated that M1 tDCS can accelerate chunk formation in early motor learning within the first training session [40]. The chunk formation process can be thought of as a transition from high uncertainty to low uncertainty in the execution of a motor skill that leads to sequence accuracy increases as the transition occurs [41]. In early stages of sequence learning with internally cued movements, efficient learning appears to prioritize learning of spatial elements and then shift to temporal components [41]. Our results support this pattern for externally cued sequences as well, as evident by the quick initial improvements in KER and the similar learning curve in both groups (Fig. 4 C). Whether timing-based performance differences resulted from enhanced encoding of temporal sequence elements across the motor network or enhanced local M1 circuitry (enlarged representation or improved activity pattern), the behavioral output that increased PI scores resulted from M1 tDCS, which supports the important role of M1 for fast motor learning and as a sequence learning substrate [42].

tDCS may have augmented Step Mania skill acquisition by hastening development of co-occurring spatiotemporal muscle patterns (i.e. ‘synergies’, [43]) for coarticulations [44] and anticipatory finger movements [45], that facilitate improved transitions between taps. Transition improvements arise as sequence elements are chunked [46] and are believed to represent temporal control optimization of sequence movements [47–49]. Due to the order and quick succession of externally paced cues with the irregular key layout, coordinated finger motions either preparing to press the next cue or making a subsequent key accessible while tapping another were beneficial for achieving optimal timing across subsequences. Coarticulation is the tendency for one element of a movement sequence to be generated in a manner facilitating other movements needed for the preceding or subsequent elements [44]. Therefore, M1 tDCS accelerating the emergence of coarticulations that facilitated element-to-element transitions is a plausible explanation for our findings. Furthermore, the timescale of our results is consistent with that of tDCS-associated synergy learning advantages that emerged primarily within session [43].

Timing is not often a tightly controlled parameter of tasks in most sequence learning studies [50] despite timing [51] and rhythm [52] being critically important to motor skill learning and performance. Relative timing is the theoretical correlate to rhythm and it refers to the relative ratio of timing components of a skill [52]. Our results suggest M1 tDCS enhanced Step Mania performance primarily through improved relative timing, which differs from previous evidence that indicated M1 tDCS affected only absolute timing [53]. However, in contrast to the present study where the context was consistent within subjects across baseline, practice, and post-test blocks, the M1 tDCS effects on timing observed by Apolinário-Souza, Romano-Silva (53) were during contextual changes for a transfer task rather than when learning was analyzed under the practiced conditions. Additionally, as complex movements may be more susceptible to tDCS accelerated learning than simpler tasks [54], the complexity of Step Mania (chunking rapid coordinated multi-finger motions; visual guidance/pre-selection [55]; visual/audio feedback action-effects [56, 57]) may be more conducive to relative timing changes to improve execution of the component finger movements versus the comparatively simpler one-finger task studied by Apolinário-Souza and colleagues [53].

Our results reinforce that M1 is an important neural substrate for motor sequence learning when the quality or outcome success of the skill relies directly upon optimized relative timing between the component movements, similar to playing a song on an instrument or shooting a jump shot, rather than the fastest possible performance of correctly ordered elements. tDCS may have impacted the neural representation of the sequence locally within M1 circuitry since it has recently been shown to have this effect in animals. In mice, increased performance after training with tDCS resulted from increased learning associated modification of specific motor circuits in M1 via enhanced correlated firing that induces Hebbian LTP within task related neural populations [58]. A similar process has been demonstrated in human auditory cortex where tDCS preferentially influenced neurons coactive with stimulation over inactive neurons [59]. On the other hand, imaging studies suggest temporal and spatial aspects of sequences may be independently encoded upstream from M1 in premotor areas [60] with only a minor component of the activity in M1 reflecting sequential characteristics [61]. During early, as compared to late, explicit sequence learning there is increased within-session coupling of M1, PM, and SMA, likely reflecting interaction functions important for fast motor learning [4, 62]. Therefore, tDCS-modulated M1 excitability may have influenced encoding of temporal aspects in premotor areas since these areas are closely reciprocally interconnected with M1 in primates [63] and Hebbian type plasticity has been demonstrated in the human M1-SMA network [64].

Successful goal-directed hand movements are highly dependent upon sensory feedback and there is extensive communication cortically between sensory and motor areas, with M1 receiving projections from many regions. [65]. Based on the apparent activity-selectivity of tDCS [58, 59, 66], we speculate that tDCS likely increased utilization of task relevant feedback by heightening sensitivity of task related M1 neural populations to converging visual and somatosensory inputs [67]. Layer II/III pyramidal cells form a broad intrinsic horizontal projection system in M1 with synaptic connections that are strengthened by LTP-like plasticity during normal motor learning [68], and within this layer are neurons that respond rapidly to somatosensory and visual feedback that converges in M1 [69–71]. Furthermore, many reciprocal interconnections between M1 and the premotor areas exist within this same layer II/III with no apparent bias in information flow [63]. Therefore, tDCS may have exacerbated the process of task related somatosensory and visual feedback shaping the neural representation in M1 locally and/or, by way of enhanced M1 activity patterns, facilitated encoding of the temporal features of the sequence in premotor areas. Considering the timing between the sequence elements was conveyed visually by the vertical distance between the scrolling arrows (Fig. 1B) and performance depended on relative timing between motions of multiple fingers, tDCS enhancing visual and somatosensory input integration in M1 is an appealing explanatory mechanism as it would be consistent with both the activity-selectivity hypothesis within M1 and the interrelated roles of premotor areas and M1 in sequence learning.

Task difficulty and baseline ability can factor into the likelihood and potential magnitude of tDCS learning effects [72], with tDCS usually benefitting novices/low performers most [26, 73]. Since our participants were naïve to rhythm games and played with their non-dominant hand, our results further support tDCS efficacy in enhancing novice skill acquisition. However, despite similar naiveté, a complex task may be easy for one person but difficult for another. This was observed while pilot testing Step Mania under normal conditions where high familiarization aptitude led to rapid performance plateaus during practice and low aptitude caused too much difficulty that rendered practice ineffective and impeded learning. M1 tDCS can improve visuomotor learning [74], but benefits may be optimized for moderately difficult (set by speed) tasks as opposed to high or low difficulty [75]. Taking this into consideration, we increased or decreased the visual cue scrolling speed for high and low baseline aptitudes, respectively, which shifted *relative* difficulty towards moderate and minimized interference from floor and ceiling effects on the emergence of tDCS effects. Therefore, task novelty notwithstanding, our results suggest that tDCS learning benefits may still be had by more experienced/skilled individuals, provided the practiced task demands moderately exceed those already managed by existing skill.

To our knowledge this is only the second study demonstrating M1 tDCS can enhance acquisition of video gaming motor skill [26], and the first specifically within the rhythm gaming genre. Video gaming inputs (i.e. controllers, keyboard and mouse) require dexterous hand and gross arm skills that, when combined with the complex visual processing and attentional demands of video games, can create motor learning paradigms to study tDCS effects with greater generalizability than traditional paradigms [26]. With the emergence of gaming as a lucrative competitive endeavor (e.g. eSports, streaming) and participants training and attempting to gain a competitive advantage, tDCS has reportedly been used to improve gaming skill [31]. Due to growing interest in such applications, commercial availability of tDCS has increased (i.e. foc.us or Halo devices); however, which regions are ideal targets for effective enhancement of specific aspects of gaming is unclear. Our results support and extend the evidence that M1 tDCS can improve gaming based motor skills. Additionally, our results also provide further indication that M1 tDCS may be able to enhance rehabilitation motor outcomes, especially given the growing interest in video game-based rehabilitation interventions and their use in conjunction with tDCS [76].

## Conclusion

tDCS shows promise as a motor learning adjuvant in many settings, but due to the task specific nature of tDCS effects on simplified laboratory tasks, there is little consensus on ideal stimulation targets to enhance motor learning for more complex real-world tasks. We have demonstrated that, compared to SHAM stimulation, M1 a-tDCS can significantly improve acquisition and performance of a complex dexterous video game skill within a single practice session. In contrast to simpler sequence learning tasks frequently used to study tDCS effects, Step Mania better represents real world skills where timing between component motions and processing incoming information relevant to task success are pivotal for learning and performance. In light of recent work highlighting that M1 might not be the primary site for plasticity that supports learning in traditional lab-based sequence tasks [61, 77], and that lab-based tasks may not probe learning a skilled continuous sequential action with high fidelity [78], the observed effects in the present study provide interesting evidence that supports the potential of M1 a-tDCS for practical real-world applications.

## Notes

### Competing Interest Statement

The authors have declared no competing interest.

